# Recovery of high-qualitied Genomes from a deep-inland Salt Lake Using BASALT

**DOI:** 10.1101/2021.03.05.434042

**Authors:** Ke Yu, Zhiguang Qiu, Rong Mu, Xuejiao Qiao, Liyu Zhang, Chun-Ang Lian, Chunfang Deng, Yang Wu, Zheng Xu, Bing Li, Baozhu Pan, Yunzeng Zhang, Lu Fan, Yong-xin Liu, Huiluo Cao, Tao Jin, Baowei Chen, Fan Wang, Yan Yan, Luhua Xie, Lijie Zhou, Shan Yi, Song Chi, Chuanlun Zhang, Tong Zhang, Weiqin Zhuang

**Author notes:** Contact Information: Name: Ke Yu, Address: Shenzhen Graduate School, Peking University, Lishui Road, Nanshan district, Shenzhen (Ke Yu).

## Abstract

Metagenomic binning enables the in-depth characterization of microorganisms. To improve the resolution and efficiency of metagenomic binning, BASALT (Binning Across a Series of AssembLies Toolkit), a novel binning toolkit was present in this study, which recovers, compares and optimizes metagenomic assembled genomes (MAGs) across a series of assemblies from short-read, long-read or hybrid strategies. BASALT incorporates self-designed algorithms which automates the separation of redundant bins, elongate and refine best bins and improve contiguity. Evaluation using mock communities revealed that BASALT auto-binning obtained up to 51% more number of MAGs with up to 10 times better MAG quality from microbial community at low (132 genomes) and medium (596 genomes) complexity, compared to other binners such as DASTool, VAMB and metaWRAP. Using BASALT, a case-study analysis of a Salt Lake sediment microbial community from northwest arid region of China was performed, resulting in 426 non-redundant MAGs, including 352 and 69 bacterial and archaeal MAGs which could not be assigned to any known species from GTDB (ANI < 95%), respectively. In addition, two Lokiarchaeotal MAGs that belong to superphylum Asgardarchaeota were observed from Salt Lake sediment samples. This is the first time that candidate species from phylum Lokiarchaeota was found in the arid and deep-inland environment, filling the current knowledge gap of earth microbiome. Overall, BASALT is proven to be a robust toolkit for metagenomic binning, and more importantly, expand the Tree of Life.

## Introduction

Metagenomics analyses accommodated numerous of approaches exploring microbial diversities, biosynthetic potentials and evolutionary relationships of earth’s microbiome (Tyson et al. 2004, Temperton and Giovannoni 2012). Specifically, the development of sequencing technologies, computational capacities and bioinformatic tools enabled genome-scale analyses, which freed our cognition of microorganisms from only cultivated isolates and significantly boosted our understandings on uncultivable microorganisms (Parks et al. 2017, Pasolli et al. 2019). Genome-resolved metagenomics was firstly applied in 2004 from low microbial diversity environment (Tyson et al. 2004), followed by a series of initiations such as Earth Microbiome Project (EMP) and the European Nucleotide Archive (ENA) which enabled us to unravel the microorganisms on a global scale (Thompson et al. 2017). For example, the latest report from EMP projects revealed more than 52,000 Metagenomic Assembled Genomes (MAGs) on species level (Amid et al. 2020, Nayfach et al. 2020), consisting a wide range of samples from environments with medium to high level of microbial complexities, such as human (Pasolli et al. 2019, Almeida et al. 2021), freshwater (Ali et al. 2020), marine (Tully et al. 2018, Reji et al. 2020), engineered environment (Ransom-Jones et al. 2017, Liang et al. 2020) and soil (Kroeger et al. 2018, Nascimento Lemos et al. 2020), etc. Such studies implementing genome-resolved metagenomic approaches have largely expanded branches of microorganisms on tree of life (Hug et al. 2016). However, despite these findings, a vast majority of microorganisms remain obscured due to 1) limitation of bioinformatic tools such as assembly and binning; 2) large unexplored area/regions with specific environmental conditions; and 3) advanced cultivation methods to be developed.

Aside from the developing innovations of culturing new microorganisms (Lewis et al. 2020), major discovery of novel species was based on sequencing-based analyses. However, major impediments that hampered us from obtaining comprehensive and high-qualitied MAGs from existing sequencing datasets are assembly and binning steps (Nayfach et al. 2020). Due to the nature of next-generation sequencing, a series of errors may occur in binning sourced from assembled short-read sequences, such as mis-clustering contigs into bins, mis-separating contigs from one genome into multiple bins, and mis-separating multiple genomes into bins sharing partial genomic sequences (Rinke et al. 2013, Yu et al. 2018, Wang et al. 2019), resulting in redundant and artificial bins that interfere the actual binning result. While the development of third generation sequencing can significantly increase the length of sequencing reads to mitigate binning errors, higher costs (e.g. Pacific Biosciences sequencing, PacBio) or error reads (e.g. Oxford Nanopore Technology, ONT) are still hindrances from acquiring intact microbial genomes with higher completeness and lower contamination (Laver et al. 2015, Wang et al. 2015). To date, the development of assembly (Bankevich et al. 2012, Peng et al. 2012, Li et al. 2015), binning (Alneberg et al. 2013, Wu et al. 2016, Kang et al. 2019, Nissen et al. 2021) and refinement tools (Song and Thomas 2017, Uritskiy et al. 2018) have allowed us recovering MAGs from relatively high diversity environments (Parks et al. 2017, Sczyrba et al. 2017), but an average of 70% of sequencing reads were still unable to be exploited on genome-resolved analyses, which proportion is even higher in more complexed environment such as soil (Howe et al. 2014, Nayfach et al. 2020). Although a growing number of bioinformatic tools have implemented algorithms on third generation sequencing datasets, such as hybrid assemblers (e.g. Unicycler, OPERA-MS, Wick et al. 2017, Bertrand et al. 2019), a systematic estimation of these tools on complexed environmental samples is crucially needed. Moreover, no study has utilized third generation sequences on post-binning approaches calibrating assembled bins and complementing genome gaps, which can maximize the exploitation of long-reads data to elevate recovered genome qualities. Therefore, to further expand tree of life, more robust assembly and binning workflows, as well as post-binning refinement methods are yet to be developed.

In addition to the demands of advanced bioinformatic tools, more investigations on remote areas/regions, especially the ones from unique environmental conditions that absent from global projects (Nayfach et al. 2020, Almeida et al. 2021) can largely expand our knowledge on earth genomic pools. As such environmental parameters (physical, chemical and biological) have direct and indirect effect on microbial communities (Berdjeb et al. 2011, Nishiyama et al. 2018, Easson and Lopez 2019, Glasl et al. 2019), exploring uncommon environment can expand the breadth of our understanding on microbial diversity and potential functions. A few studies have already launched targeting rarely explored environments on deserts (Finstad et al. 2017), oil fields (Eze et al. 2020), hydrothermal vents (Anderson et al. 2017) and non-marine soda lakes (Vavourakis et al. 2018), but such number of investigations were still scarce to support the expansion of microbial diversity, compare to a vast majority of studies on marine, soil and human-associated microbiomes (Hoshino et al. 2020, Nascimento Lemos et al. 2020, Almeida et al. 2021).

In this study, we introduce a highly robust toolkit BASALT (Binning Across a Series of AssembLies Toolkit) to robustly recover microbial genomes from sequencing datasets using a series of innovative methodologies for assembly, binning and refinements. Firstly, BASALT uses high-throughput assembly methods to automatically assemble/co-assemble multiple files in parallel to reduce the manual input; Next, BASALT incorporates self-designed algorithms which automates the separation of redundant bins to elongate and refine best bins and improve contiguity; Further, BASALT facilitates state-of-art refinement tools using third-generation sequencing data to calibrate assembled bins and complement genome gaps that unable to be recalled from bins; Lastly, BASALT is an open frame toolkit that allows multiple integration of bioinformatic tools, which can optimize a wide range of datasets from various of assembly and binning software. Using BASALT, we performed a case study on sediment samples of Aiding Lake, Xinjiang China, a deep-inland hypersaline lake with high aridity in the surrounding area (Figure S1, Guan et al. 2020), which is different from common chlorite saline aquatic system. Based on the superior number and quality of MAGs obtained via BASALT, we present taxonomic and genetic profiles on prokaryotic microbial communities of lake sediment samples, including two Lokiarchaeota species from a recently discovered archaeal superphylum Asgardarchaeota (Zaremba-Niedzwiedzka et al. 2017) that also found in our samples. Here, we highlighted that BASALT can efficiently enhance the assembly and binning processes on metagenomic sequences, which allows in-depth investigations on the existing dataset by increasing overall and individual MAG quality. By acquiring more high-qualitied MAGs, we could potentially unfold much more knowledges on known or unknown microbial taxa, potential functions and host-microbial interactions, which further expand the tree of life.

## Results

### Environment, pipeline and availability

BASALT is command line software based on Python scripts with a series of modules, each containing one or more algorithms/programs addressing data processing or analysis. Overall, BASALT is an automated program running with one command line, while a few checkpoints are set in each module to accommodate users to customize their preference to start at any checkpoint as needed. Further details of outlines and algorithms are available at (https://github.com/EMBL-PKU/BASALT).

### BASALT workflow

BASALT is a versatile toolkit that provides comprehensive pipeline from mapping, automated binning to post-binning refinement that enable users to retrieve non-redundant and high-qualitied MAGs from metagenome samples. Overall, BASALT contains four major modules including Automated Binning, Bin Dereplication, Bin Refinement and Reassembly, where five core algorithms were implemented in these modules including Core Contigs Identification, Bin depth normalization, Outlier Removal, Contigs Retrieval and Restrained Overlap-Layout-Consensus (rOLC, Figure 1). As BASALT enables multiple assembly or co-assembly datasets from short-read sequences (next-generation sequencing, NGS) or long-read sequences (third-generation sequencing, TGS) as input files, it is expected that a potential increase of reads utilization is available (Stewart et al. 2018). By importing multiple sequences with coverage information, BASALT conducted automated binning using prominent tools such as MetaBAT2, Maxbin2 and CONCOCT (Alneberg et al. 2013, Wu et al. 2016, Kang et al. 2019) that generated hybrid bin-sets. Raw hybrid bin-sets were firstly filtered with a homebrew Bin Dereplication module to remove replicated bins obtained from the same assembly. The non-redundant bins of each assembly were then merged and categorized into different groups at a customized average nucleotide identity (ANI) cutoff. Each group of bins were further filtered by the homebrew algorithm which classifies core contigs that further enables identification of redundant bins before selecting into a single, hybrid bin-set obtained from all samples. The selected bin-set was further filtered using a critical Outlier Removal algorithm, which integrated coverage and tetranucleotide frequency (TNF) to remove outliers by using an interquartile ranges (IQR) method with multiple thresholds. Next, sequences from assembly files were retrieved by connecting with existing contigs in the OR-filtered bins to create an expanded sequences pool with potential connected contigs, while BASALT compared and selected the refined bins with higher quality value. Further, a restrained Overlap-Layout-Consensus (rOLC) step was conducted to overlap the replicated bins obtained from different assemblies into OLC-merged bins, followed by the reassembly step to generate the finalized bin-set. Notably, the sequence retrieval and reassembly step allowed utilization of long-read sequences obtained from TGS to complement gaps and join overlapped regions on the bins to increase completeness and reduce contamination.

**Figure 1.**
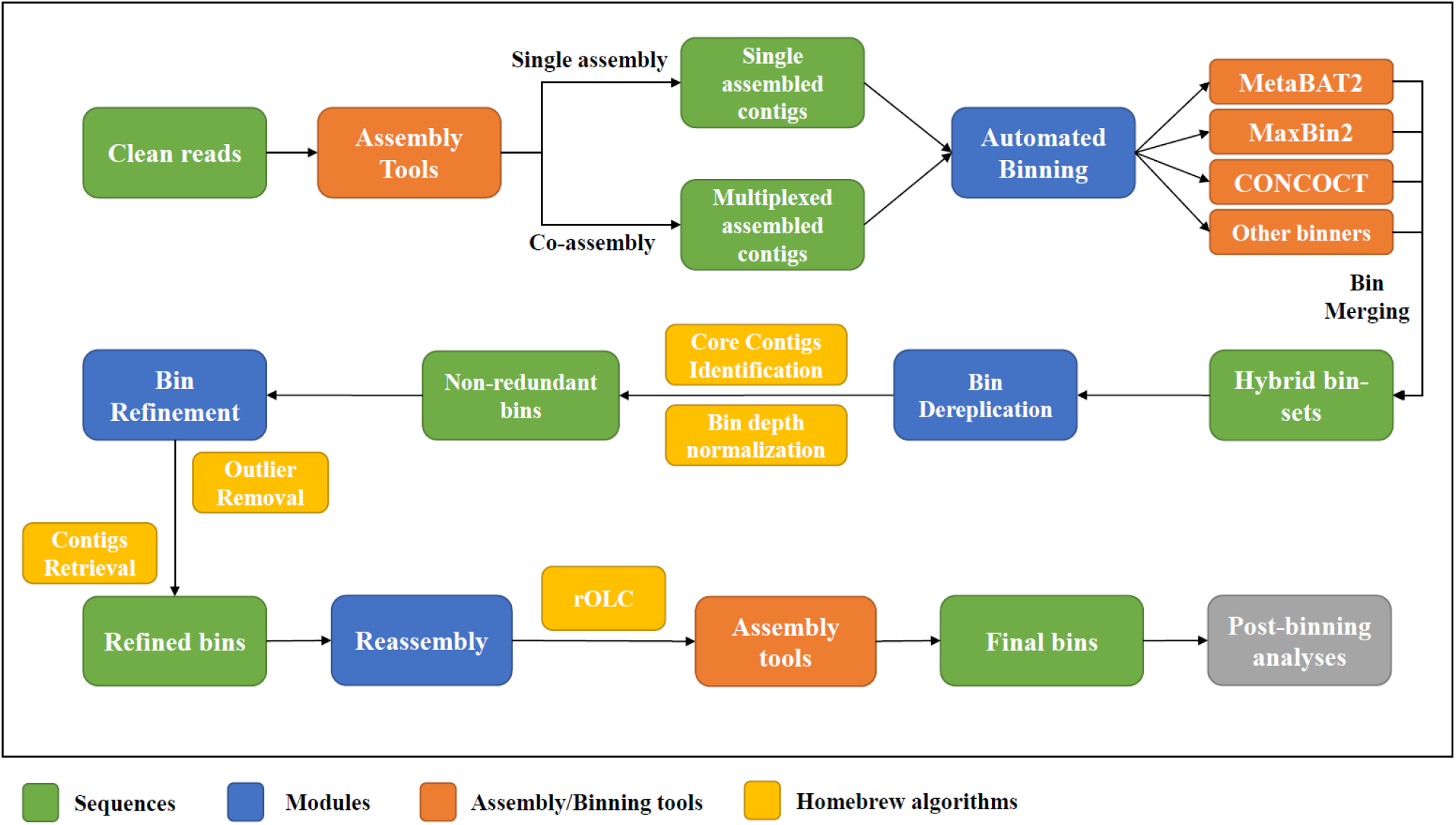
BASALT workflow for assembly, binning and refinement of metagenomic sequencing data (blocks in green). BASALT contains four major modules (blocks in blue) including Automated Binning, Bin Dereplication, Bin Refinement and Reassembly, where five core algorithms (blocks in yellow) were implemented in these modules including Core Contigs Identification, Bin depth normalization, Outlier Removal, Contigs Retrieval and Restrained Overlap-Layout-Consensus (rOLC). In addition to core workflow, assembly and binning tools (blocks in orange) were also flexibly embedded in BASALT.

### BASALT improves recognition of non-redundant bins

BASALT pipeline enables multiple input files for assembly, including long-reads files for hybrid assembly. The advantage of input with multiple files is not limited to the reduction of computational time but could also generate more bins than individual assembled samples. For example, binning using multiplexed samples generated 16.3%, 14.2% and 11.1% more non-redundant MAGs when using DAS Tool (MCM), VAMB and metaWRAP (MCM), respectively on CAMI-medium dataset (Table S1, Figure S2), while the increasing rate on CAMI-high dataset were 8.2%, 11.0% and 9.0%, respectively (Table S1, Figure 2A). Despite the advantage above, a major drawback using multiplexed samples was the byproducts of replicated bins, which were considered as redundant or pseudo- genomes. To address this issue, BASALT pipeline accommodates a Bin Dereplication module with homebrew algorithms which can remove redundant bins generated from the multiplexed assemblies as well as hybrid bin-merging from automated binning, resulting in optimized, non-redundant bin-sets (Figure 1). Comparing with standard CAMI-medium and -high genomes, DAS Tool, VAMB and metaWRAP generated 85.9%, 77.1%, 95.0% (CAMI-medium) and 84.8%, 52.3%, 72.7% (CAMI-high) redundant MAGs, respectively (Figure 2A, Figure S2), while no redundancy was observed in BASALT MAGs, suggesting high redundancy rate were existed using the first three binning tools/toolkits. Remarkably, BASALT Bin Dereplication, Refinement and Reassembly modules can not only eliminate redundant MAGs generated from DAS Tool, VAMB and metaWRAP pipelines, but also increase the overall quality of non-redundant MAGs (Co-assembled data refined with BASALT, CAB, Figure 2A), suggesting good efficiently of redundancy removal and quality improvement using BASALT from co-assembled bins.

**Figure 2.**
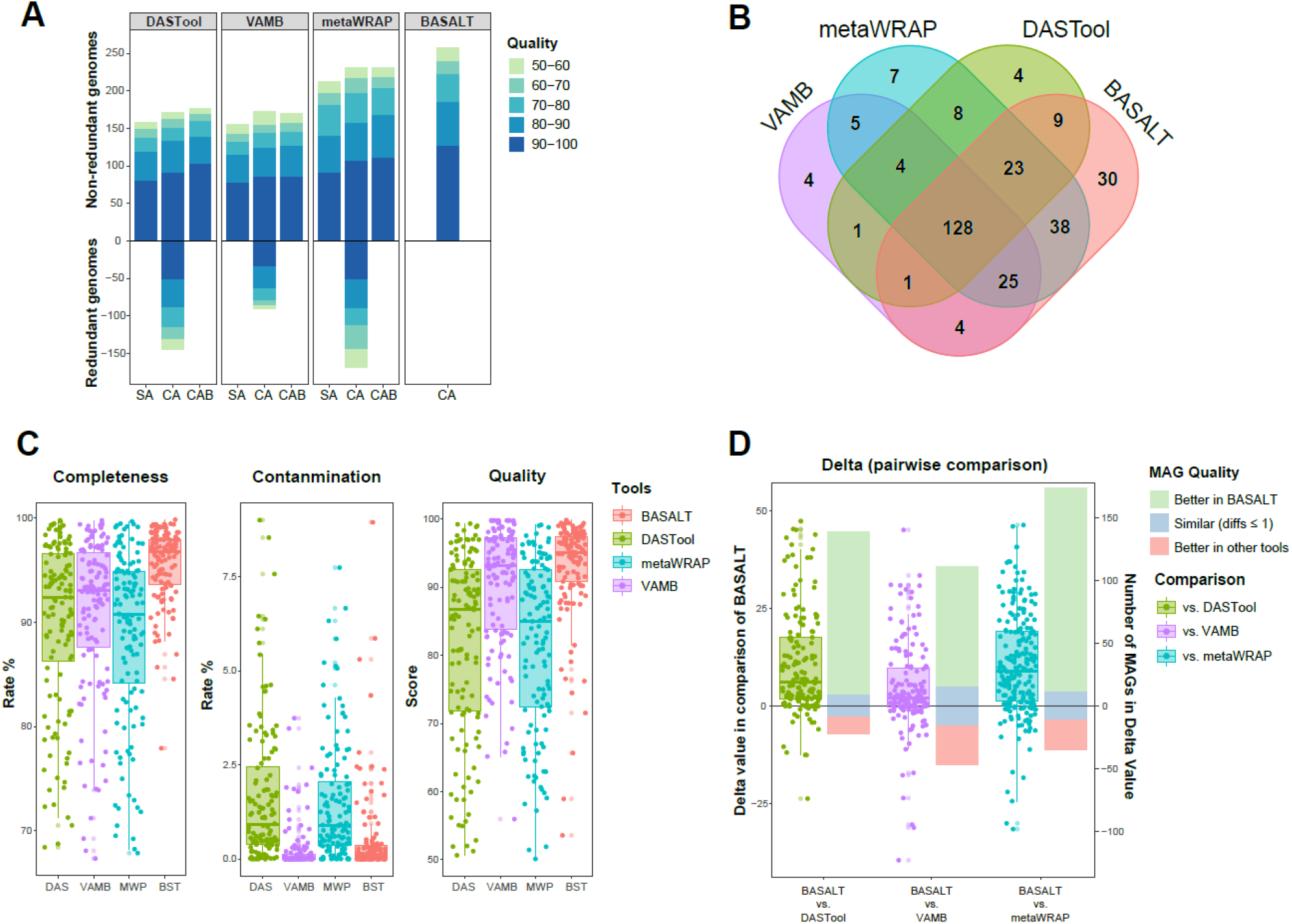
Comparison of BASALT with other binning tools/pipelines on CAMI-high dataset. **A)** Number of MAGs recovered from CAMI-high dataset using DASTool (MaxBin2, CONCOCT and MetaBAT2, MCM), VAMB, metaWRAP (MCM) and BASALT. In the first three tools, Co-assembly (CA) resulted in higher number of non-redundant MAGs compare to single assembly (SA) approach, while BASALT refinement module (Co-assembly refined with BASALT, CAB) removed redundant MAGs and generated higher quality of MAGs in the co-assembly approach. Color of bars indicated the quality of MAGs (50-100, from light to dark). **B)** Venn diagram showing number of MAGs recovered using different tools. There were 128 MAGs found shared across all tools, while 4, 4, 7 and 30 MAGs were uniquely recovered using DAS Tool (green), VAMB (purple), metaWRAP (cyan) and BASALT (red) pipelines, respectively. **C)** Completeness, contamination and quality of 128 shared MAGs recovered using DAS Tool (DAS, green), VAMB (purple), metaWRAP (MWP, cyan) and BASALT (BST, red). MAGs recovered using BASALT had lower contamination rate compared to DAS Tool and metaWRAP, and higher completeness and quality compared to all other tools. **D)** Pairwise comparison of MAGs shared by corresponding pairs of tools. Overall, BASALT was superior in obtaining higher qualitied MAGs compared to other tools. The number of MAGs that BASALT gained higher quality value (bars in light green) was much more than the number of MAGs that other tools gained higher quality value (bars in light red) or had similar quality value (difference of value ≤ 1, bars in light blue).

### BASALT generates higher number and qualitied MAGs in synthetic microbial communities

The BASALT refinement module not only removes redundant bins generated in other pipelines, but also improves the quality and number of MAGs. In comparison of MAGs (Quality score ≥ 50) obtained via different toolkits, BASALT resulted in 27.1%, 7.0%, 1.7% (CAMI-medium) and 50.9%, 50%, 11.7% (CAMI-high) more non-redundant MAGs than DAS Tool, VAMB and metaWRAP, respectively (Figure 2A, Figure S2, Table S1). Moreover, in top-qualitied CAMI-high MAGs (Quality score ≥ 90), BASALT obtained 38%, 47.7% and 17.6% more non-redundant MAGs than DAS Tool, VAMB and metaWRAP, respectively (Figure 2A). This result suggested BASALT is more robust retrieving high-qualitied and non-redundant MAGs from metagenome samples, especially on those samples with higher complexity.

To individually evaluate the BASALT refinement module, a further refinement step was performed on CAMI-high datasets processed with DAS Tool, VAMB and metaWRAP. Although less MAGs were recovered compare to the result using comprehensive BASALT pipeline, 3.7%, 2.4% and 7.0% more MAGs (Quality score ≥ 80) were retrieved, respectively, compared to the default approaches of other toolkits (Table S1, Figure 2A). This result suggested that BASALT can also optimize number and quality of MAGs even based on the datasets processed with other pipelines.

Comparing MAGs recovered from the abovementioned toolkits with reference genomes in CAMI-high dataset, 128 MAGs were found universally presented across all pipelines, while 4, 4, 7 and 30 MAGs were uniquely recovered using DAS Tool, VAMB, metaWRAP and BASALT pipelines, respectively (Figure 2B). In comparison of the 128 shared MAGs in their completeness, contamination and quality score (completeness – 5*contamination) across different toolkits, BASALT has the overall advantages in acquiring high-quality and low contamination MAGs in comparison with others (Figure 2C). Further, we performed pairwise comparison between BASALT and the other tools of shared MAGs and calculated delta value (difference of MAG quality between two tools) on the same genome. The delta value showed BASALT could retrieve 10, 3.1 and 6.8 times of high-quality MAGs compared to DAS Tool, VAMB and metaWRAP, respectively (Figure 2D), indicating that BASALT can substantially obtain better quality of MAGs comparing with other tools. In summary, BASALT can retrieve a greater number of non-redundant MAGs from medium-high complexed samples with higher qualities.

### BASALT efficiently retrieves genomes from high complexity samples

We performed four metagenome samples from Aiding Lake sediments using both BASALT to evaluate the capacity of BASALT on improving genome qualities recovered from complexed environmental samples. Using BASALT, assembled bins were quality-checked with CheckM (Parks et al. 2015), resulting in 426 non-redundant MAGs (completeness – 5*contamination ≥ 50, mean completeness = 79%, mean contamination = 1.7%, mean quality value = 70.4), including 113 MAGs above high-quality level (quality ≥ 80). As a majority of metagenomic sequences could not be utilized to recover high-qualitied genomes (Nayfach et al. 2016, Nayfach et al. 2020), we estimated the efficiency of sequences utilized in the lake sediment samples, resulting in 26.9% of reads mapped to the MAGs, which largely improved the utilization rate of metagenomic sequences in recovered MAGs compared to other samples with high complexity (Howe et al. 2014).

In reference to Genome Taxonomy Database (GTDB, Parks et al. 2018), all 426 MAGs from BASALT pipelines were annotated spanning 39 bacterial and 8 archaeal phyla. MAGs classified into bacterial phyla were mainly focused in Patescibacteria, Chloroflexota, Verrucomicrobiota, Bacterioidota, Proteobacteria and Desulfobacterota, while MAGs classified into archaeal phyla were Halobacteriota, Thermoplasmatota, Asgardarchaeota, Thermoproteota, Iainarchaeota and Micrarchaeota in archaeal domain (Figure 3A), representing a unique characteristic of microbial communities in the Salt Lake sediment. Among these 426 MAGs, 98.9% bacterial MAGs and 100% archaeal MAGs could not be assigned to any known species from GTDB (ANI < 95%). At genus level, 57.6% bacterial MAGs and 66.7% archaeal MAGs could not be assigned to any known genus from GTDB. This result suggested that there was a large repository of genetic pool to be uncovered in Salt Lake sediments.

**Figure 3.**
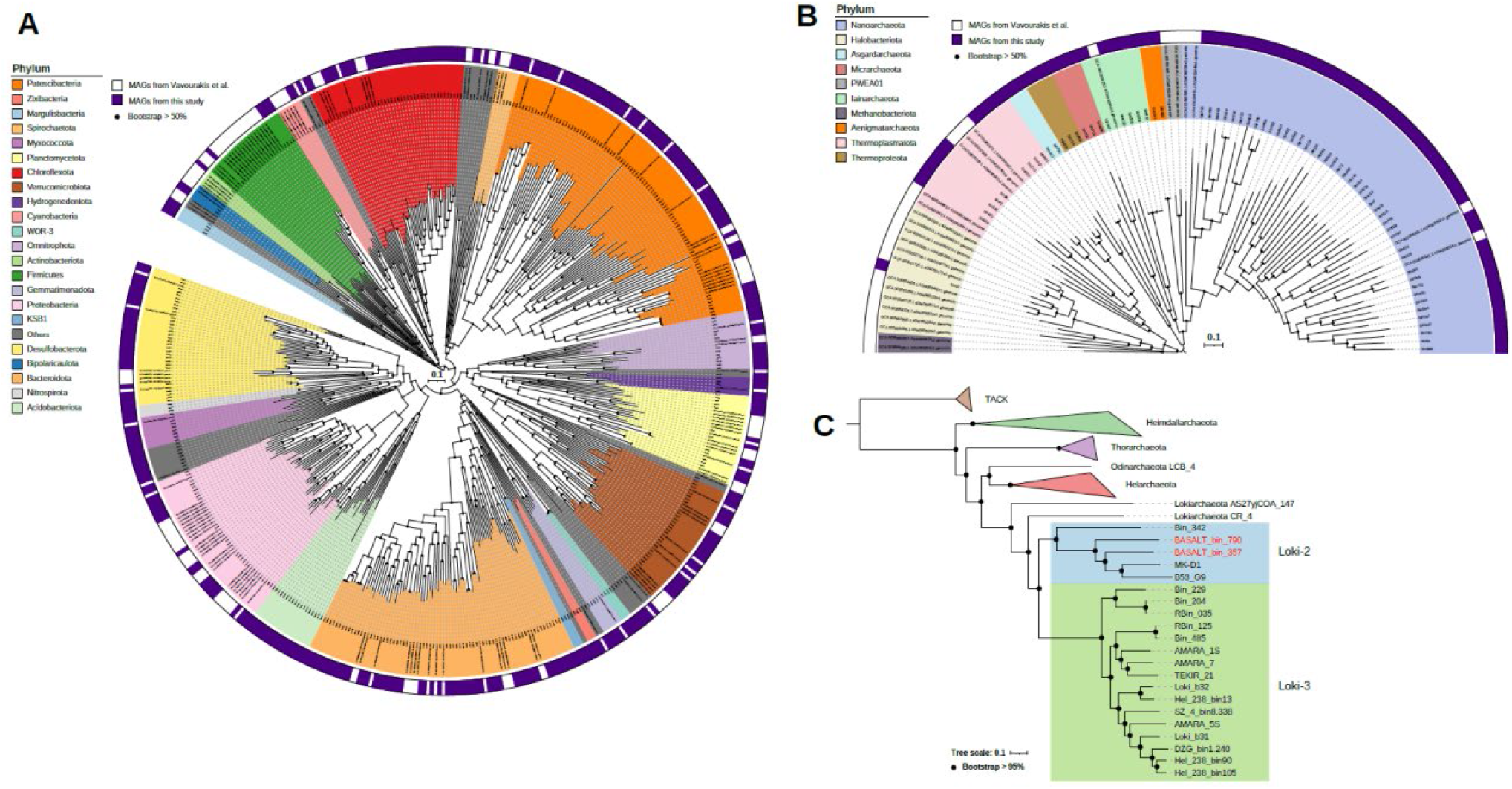
Phylogenetic trees of **A)** bacterial MAGs from Vavourakis et al. (2018) and this study, **B)** archaeal MAGs from Vavourakis et al. (2018) and this study, and **C)** Maximum-likelihood tree of Asgardarchaeotal MAGs based on protein encoded genes from the whole genome. The unrooted phylogenetic trees in A) and B) were constructed with Fasttree, while the Maximum-likelihood tree in C) was constructed with PHYML and re-rooted with superphylum TACK. The two Lokiarchaeota MAGs found in this study were highlighted with red in C), and all genomes used to create trees that not obtained from this study were listed in Table S2.

To further validate the MAGs recovered from Aiding Lake sediments, we compared these assembled genomes with another study by Vavourakis et al. (2018) where samples were collected from several soda lakes in the Kulunda Steppe (south-western Siberia, Altai, Russia). Generally, the two datasets shared a vast majority of phyla (Figure 3A), indicating that despite the different geographical location, bacterial assemblages of the Salt Lakes might be similar. Specifically, diverse bacterial taxa from phylum Patescibacteria were observed in our study, which paralleled with the study by Vavourakis et al. (2018). On the other hand, in the unique of phyla observed in two datasets, Acidobacteriota, Nitrospirota, Zixibacteria, WOR-3 and KSB31 were only present in this study (Figure 3A). In archaeal communities, MAGs from phyla Halobacteriota, Thermophasmatota, Iainarchaeota and Nanoarchaeota were observed in both datasets, while phyla Methanobacteriota and PWEA01 were only present in Vavourakis et al. (2018) and phyla/superphyla Asgardarchaeota, Thermoproteota, Micrarchaeota and Aenigmatarchaeota were only present in our study (Figure 3B). In the shared phyla, a large number of MAGs from phylum Nanoarchaeota found, but only one MAG from phylum Halobacteriota observed in this study, while a large community of Halobacteriota was observed in Vavourakis et al. (2018). The difference of the two datasets might be due to the physiochemical parameters between two contrasting sites of lakes (Sorokin et al. 2014), but such differences could also be related to the different strategies of assembly, as well as binning pipelines.

### Newly discovered Lokiarchaeota species from a deep-inland non-marine hypersaline lake

Using four metagenome samples, BASALT pipeline expanded 421 of known species on prokaryotic phylogenetic tree with samples from one Salt Lake. Remarkably, two MAGs classified as Lokiarchaeota belongs to the superphylum Asgardarchaeota, were also observed in our results. Although previous study has reported Asgardarchaeota phyla found in hypersaline lakes (Bulzu et al. 2019), to the best of our knowledge, this is the first time that archaeal phylum Lokiarchaeota was found from the deep inland non-marine samples. Comparing with other MAGs/isolates obtained in previous studies, the two MAGs found in our study were grouped with Loki-2 MAGs/isolates (Figure 3C), next to *Ca. Prometheoarchaeum syntrophicum*, a strain isolated from a deep-sea sediment sample at Nankai Trough, Japan (Imachi et al. 2020). Interestingly, other Loki-2 MAGs were found globally, including Bin_342 from Shark Bay, Australia (Wong et al. 2020) and B53_G9 from Guaymas Basin, US (Seitz et al. 2019), suggesting that Loki-2 species were widely distributed not only in marine sediments but also in the deep-inland terrestrial environment.

## Discussion

BASALT is superior in quality and number of MAGs on low (132 genomes) to medium (596 genomes) complexity samples. In regards of MAG quality assessment, CheckM (Parks et al. 2015) is widely used in a vast majority of studies. However, in the presence of standard CAMI datasets, we calculated the MAG quality against the corresponding genomes that can result in more accurate evaluations. Notably, our dereplication module implemented in the integration step can efficiently remove redundant bins that generated in co-assembly and bin selection steps, which was evidenced in the test analysis using both CAMI-medium and CAMI-high datasets (Figure 2A). Although there are other redundancy removal methods such as dRep (Olm et al. 2017), result suggested that redundant genomes cannot be efficiently identified and removed using dRep on CAMI-medium or CAMI-high datasets (Table S1), suggesting that BASALT Bin Dereplication gains more advantages in removing redundant bins under higher complexed samples. Due to the scarcity of standard synthesized community with high complexity to date, further works testing the efficiency of Bin Dereplication module on the standardized samples with high complexity are needed.

BASALT-integrated Bin Dereplication module can efficiently remove redundant bins generated by binning tools. However, due to the major impediments of short-fragmented technology of next-generation sequencing (NGS) and post sequencing algorithm, recovered MAGs cannot specify differences of genomes at strain level with high similarity. While long read sequencing such as ONT has become more popular in the current metagenomic studies (Jain et al. 2016), high error rates still required to be rectified by NGS. In this study, third-generation sequences were innovatively integrated into our toolkit where long reads can be used to amend sequences and fill gaps on assembled genomes, which can improve the overall quality of bins and increase the number of high-qualitied MAGs. Although standard dataset with high complexity is currently not available in the database, our case study revealed that Salt Lake sediments can be considered as samples with relatively high complexity (6,993 ZOTUs from 16S rRNA gene amplicons), which was comparable with other studies on soil samples regardless the primer selection (Fulthorpe et al. 2008, Xiong et al. 2021). In the context of high complexed samples of Salt Lake sediments, BASALT could also efficiently conduct reads utilization in metagenomic binning with 26.9% mapped sequences, which to some extent helped to resolve the difficulty raised in the EMP project (Nayfach et al. 2020). Prospectively, the trend in the development of third-generation sequencing has inspired that further exploitation of long-read sequences can increase the resolution of MAGs at strain or single nucleotide polymorphism (SNP), which may boost the outcome of recovered genomes in high complexity samples. Therefore, future development should focus on the combination of NGS and TGS to improve the efficiency of binning, which consequently expand our knowledge on the tree of life.

In addition to the BASALT toolkits introduced in this study, one major finding in the case study was the discovery of two novel Lokiarchaeota genomes from the sediment samples of Aiding Lake. Lokiarchaeota belongs to a recently discovered superphylum Asgardarchaeota, which is a hot topic linked to the origin of eukaryotes (Spang et al. 2018). To date, candidate species of Lokiarchaeota were universally found near deep sea hydrothermal vents and marine sediments (Spang et al. 2015, Spang et al. 2018, Hoshino et al. 2020, Wong et al. 2020, Yin et al. 2020). A recent study has found candidate Lokiarchaeota present in hypersaline lakes near Black Sea (Bulzu et al. 2019), whereas our study highlighted the first time that Loki-2 species were found from deep-inland hypersaline lake sediments. Given the genetic analysis on Lokiarchaeota along with other candidate Asgardarchaeota species (Zaremba-Niedzwiedzka et al. 2017, Seitz et al. 2019, Imachi et al. 2020, Wong et al. 2020, Yin et al. 2020) have suggested that candidate Lokiarchaeota species were adaptive in marine environment with distinct metabolic pathway, such as lignin or protein degradations. Thus, it was unlikely that candidate Loki-2 species were newly emerged in the deep-inland hypersaline lake. Therefore, the discovery in this study might have provided a landmark that candidate Lokiarchaeota species might exist in the ancient age before the plate movement event occurred. However, insufficient MAGs/isolates revealed to date hampered us to make rigid conclusions, that more investigation on this group of archaea is critically required in the future studies.

## Methods

### Overview of BASALT

BASALT is a versatile toolkit that recovers, compares and optimizes MAGs across a series of assemblies assembled from short-read, long-read or hybrid strategies. We established five homebrew algorithms to carry out Core Contigs Identification (CCI), Bin Depth Normalization, Outlier Removal (OR), Contigs Retrieval, and restrained overlap-layout-consensus (rOLC). These algorithms consist three core modules of BASALT, such as Bin Dereplication, Bin Refinement and Bin Reassembly modules. Besides, BASALT contains an autobinning module that uses mapping tools (e.g. bowtie2, Langdon 2015) and binners (e.g. MetaBAT2, Maxbin2, and CONCOCT) to generate a raw hybrid bin-set after input of assemblies (Figure 1).

### Dereplication of redundant bins

Incompleteness and contamination of MAGs would hinder the dereplication of bins from the hybrid bin-set. We developed an effectively strategy that can remove most of the potential contaminated sequences while large number of sequences are still kept to carrying out precise redundancy identification. Different from previously reported genome-wide ANI-based and marker gene- based de-replicating methods such as dRep (Olm et al. 2017), the present method firstly generated a core contig pool by filtering out potentially contaminated contigs of target bins. The depth and tetranucleotide frequency (TNF) value of the selected contigs ranked from a range of 25 to 75 percentile of all the contigs of target bins. Then, selected bins were grouped into different raw bin-sets based on overall similarity of core contigs. Secondly, we evaluated the sequencing depth discrepancy among bins across different bin-sets to ensure the correct identification of redundant bins with relative low similarity. The average depth of one bin should be equivalent to the average depth of the core contigs of this bin. For neutralizing the sequencing depth discrepancy yielded by mapping the same reads to different assemblies, we designed a method to calculate the normalization ratio between two possibly redundant bins, which identifies near-identical sequences (99.8% similarity across longer than 50% of the whole length of sequence) between two potential redundant bins as candidate bins. A depth normalization ratio between candidate bins was then calculated by using the depth of these contigs, which was then used to neutralize the average depth of candidate bins. Those candidate bins with similar average sequencing depth (delta ≤ 10%) were considered as redundant bins to be removed. Finally, the bin with better quality value (completeness – 5* contamination) estimated by CheckM (Parks et al. 2015) was kept being the better bin for further refinement and reassembly.

### Refinement

BASALT refinement module contains two adversary processes: Outlier Removal (OR) and Contig Retrieval. To effectively remove contaminated contigs from a certain bin, we designed an outlier removal algorithm that removes contigs with an outlier value of sequencing depth or TNF based on an interquartile range (Formula 1), while different thresholds (k) were set (e.g. 1, 1.5, 3) to determine these contigs. In the context of bin quality, OR keeps bins with higher quality value, while bins with lower quality were discarded. If no refined bin with higher quality than the original bin was generated, OR would acquiescently eliminate sequences marked as depth or TNF outliers under the threshold of 3. Notably, the default setting of OR mainly removes contaminated sequences, but it may cause an unnecessary removal of contigs due to restricted threshold.

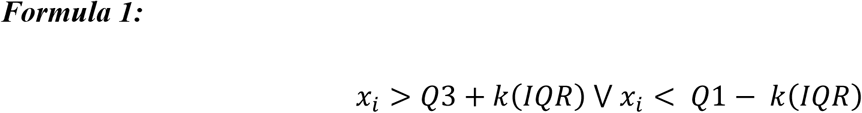

In this formula, Q1 and Q3 stands for 25 and 75 percentiles of all contigs, where IQR (interquartile ranges) was calculated by Q3 – Q1. k ≥ 0.

Contig Retrieval algorithm was designed to retrieve sequences that have not been clustered into the target bin in the binning process, especially multicopy sequences or unnecessarily removed sequences by Outlier Removal. Contig Retrieval identifies contigs that potentially connected to existing sequences in the target bin. These candidate contigs were further assessed by an interquartile range method to remove depth or TNF outliers as described above and connected to the target bin by paired-end tracking (Albertsen et al. 2013) or long-read mapped method, forming a refined bin. Refined bins were expected to have higher quality value than target bins which were further selected to form a refined bin-set for further reassembly.

### Reassembly

BASALT reassembly module includes a restrained overlap-layout-consensus (rOLC) process and a reassembly process. The rOLC algorithm was designed to retrieve sequence which was not included in the BASALT binning and refinement processes. Specifically, this process re-utilized sequences from redundant bins identified and removed from the dereplication module to overlap the sequences from the target bin. Using a loose threshold (overlap length: 300 bp; similarity: 99%), rOLC algorithm aggressively recorded the redundant candidate sequences removed from the target bin in the previous steps. As rOLC process may increase the contamination of sequences, a reassembly step was implemented after rOLC to amend the contamination caused by rOLC, and more importantly, to precisely elongate the length of sequences in the target bin. In the reassembly process, short-read or long-read sequences were extracted from datasets by Bowtie2 and Minimap2, respectively (Langdon 2015, Li 2018). Three state-of-art assemblers were implemented in the reassembly step: SPAdes, FLYe, and Unicycler (Bankevich et al. 2012, Wick et al. 2017, Kolmogorov et al. 2019) to carry out short-read reassembly, long-read reassembly and hybrid reassembly, respectively. A final bin selecting process was exploited to select the best bins from the reassembly step to form the final bins for post-binning analysis.

### Sample collection

Salt Lake sediment samples were collected in July 2018 from Aiding Lake, an arid region in Turpan City, Xinjiang Uygur Autonomous Region (42°52′9″ N, 89°03′5″ E). Briefly, about 50 grams of sediment samples (n = 4) were randomly collected at 0-10 cm depth in the lake into sterile 50 ml falcon tubes. Samples were immediately placed on dry ice before brought to laboratory and transferred to −80 °C freezer until further DNA extraction was performed.

### DNA extraction and sequencing

Frozen stored sediment samples (~250 mg dry weight per sample) was used to extract genomic DNA using DNeasy PowerSoil Kit (Qiagen, Hilden, Germany), following the manufacturer’s instructions. Extracted DNA was quality checked by NanoDrop 2000 (Thermo Fisher Scientific, Waltham, Massachusetts, US), quantity checked by Qubit Fluorometer (Thermo Fisher Scientific) and PCR checked to confirm the amplifiability.

For 16S rRNA gene amplicon sequencing, barcode as a marker for each sample DNA was added at the 5’ end of primers targeting V4 region for bacterial and archaeal communities (515F-806R, Caporaso et al. 2012). Sequencing was performed at MAGIGENE, Guangzhou, China on an Illumina NovaSeq 6000 platform (2 × 250 bp paired end chemistry).

For shotgun metagenomics sequencing, quantity and quality checked genomic DNA was sent to Novegene Co., Ltd, Nanjing, China on an Illumina NovaSeq (2 × 150 bp paired end chemistry).

### Sequencing processing

To evaluate the efficiency of BASALT, standard Critical Assessment of Metagenome Interpretation (CAMI) datasets including a simple-complexity (132 genomes, CAMI-medium) and a medium-complexity (596 genomes, CAMI-high) synthesized communities were downloaded from (https://data.cami-challenge.org/participate) (Sczyrba et al. 2017). Raw paired-end reads were initially filtered using fastp (Chen et al. 2018). Fifty percent of bases were filtered based on a minimum quality score of 5 and sequence length of 150 bp, allowing no ambiguous bases. Clean reads were individually and co-assembled using SPAdes (version3.14.1, Bankevich et al. 2012) into contigs specifying k-mer sizes of 21, 33, 55, 77 and finally reserved contigs > 1,000 bps. To compare with other binners/toolkits, filtered contigs were processed with DASTool, VAMB, metaWRAP and BASALT, respectively. The redundancy, completeness and contamination of the MAGs were calculated against standard CAMI datasets using a homebrew script *Bin_quality_evaluation.py* available on github (https://github.com/EMBL-PKU/BASALT) to ensure high accuracy of results obtained from the four binners/toolkits. High-quality MAGs (completeness - 5*contamination ≥ 50%) were kept for further statistical analysis, whereas bins did not meet the quality were discarded.

For Aiding Lake sediment samples, raw 16S rRNA gene amplicon sequences were processed using Quantitative Insights Into Microbial Ecology (QIIME2) pipeline (http://qiime.org) (Caporaso et al. 2010). DADA2 was used to filter low-quality sequences with lengths < 230 bp, remove chimeric sequences, singletons, and join the quality-filtered paired-end reads. Unique sequences (100% similarity) were taxonomically assigned using Naive Bayes classifier against the SILVA 16S rRNA gene reference alignment database (release 123) (Pruesse et al. 2007). To compare the complexity of Salt Lake sediment samples with other high-complexed samples such as soil, sequences were rarefied to an even sampling depth at 10,000 reads per sample, before singleton was removed from the generated OTU table. Overall, a total number of 6,993 ZOTUs (Zero-radius OTUs) were obtained.

For shotgun metagenomic sequences of Aiding Lake sediment samples, sequences were processed following the same procedure on CAMI-medium and CAMI-high datasets using BASALT. The completeness and contamination of the MAGs were then estimated using CheckM version 1.1.3 (Parks et al. 2015) with lineage-specific marker genes and default parameters, with only high-quality MAGs (completeness - 5*contamination ≥ 50%) were kept for further analyses.

### Phylogenetic analysis

The GTDB-Tk version (version1.4.1, Chaumeil et al. 2020) program was used to assign taxonomic classifications to the MAGs (release r95). To make comparison with another study of soda lake samples (Vavourakis et al. 2018), dereplicated MAGs were downloaded from NCBI Assembly database and phylogenetic analyses were conducted based on MAGs from both studies using Fasttree (Price et al. 2010). For phylogenetic analysis of Asgardarchaeota, MAGs/isolates were downloaded from other studies listed in Table S2, and Maximum-likelihood tree was constructed using PHYML version 3.0 (Guindon et al. 2010) with 1000 bootstrap iterations. Phylogenetic trees were visualized and edited in the iTOL (https://itol.embl.de) online platform (Letunic and Bork 2019).

## Code availability

All BASALT codes including homebrew scripts for quality checking against standard datasets are available at (https://github.com/EMBL-PKU/BASALT).

