## Supplementary Materials for "Recovery of high-qualitied Genomes from a deep-inland Salt Lake Using BASALT"

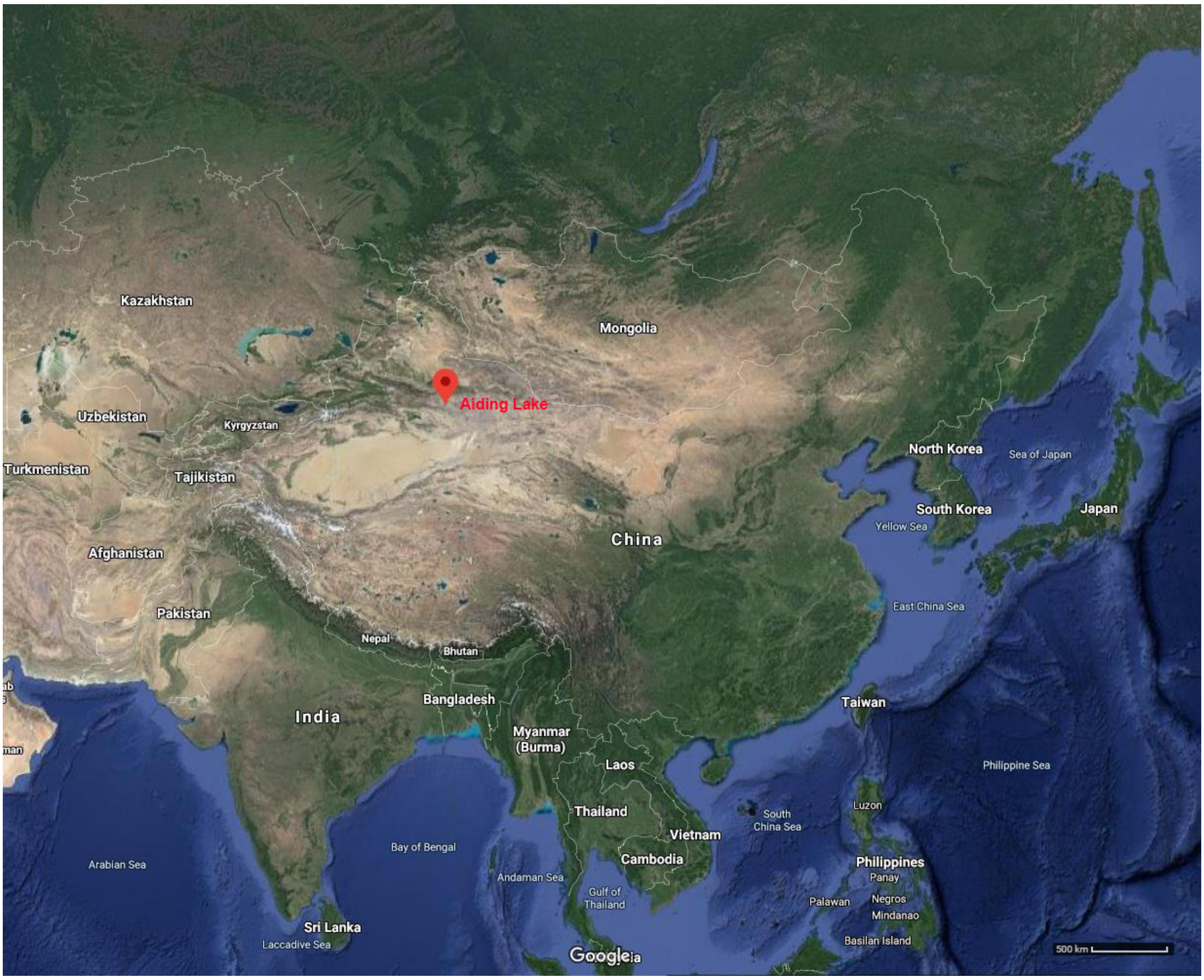


**Figure S1.** Location of Aiding Lake highlighted in the map.

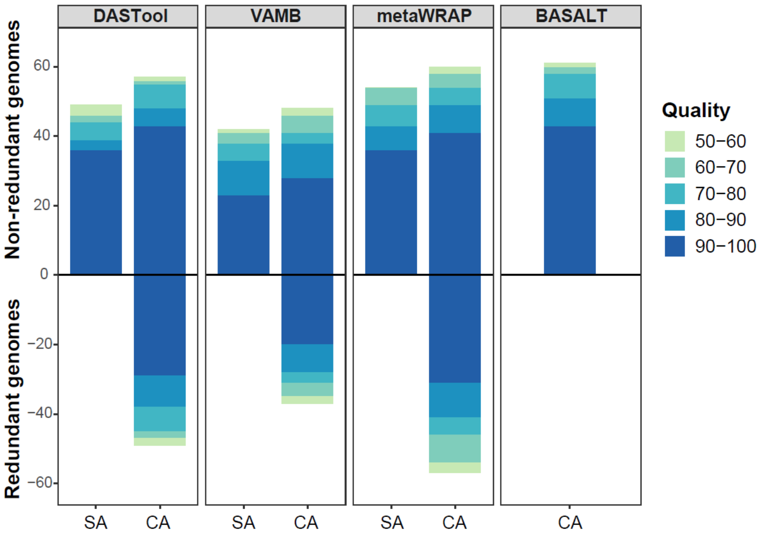


**Figure S2.** Comparison of BASALT with other binning tools/pipelines on CAMI-medium dataset. Number of MAGs recovered from CAMI-medium dataset using DASTool (MaxBin2, CONCOCT and MetaBAT2, MCM), VAMB, metaWRAP (MCM) and BASALT. In the first three tools, Co-assembly (CA) resulted in higher number of non-redundant MAGs compare to single assembly (SA) approach, while BASALT generated higher number and quality of MAGs in the co-assembly approach. Color of bars indicated the quality of MAGs (50-100, from light to dark).

**Table S1.** Summary of MAGs obtained by different toolkits/binners. Quality score was calculated by completeness (CPN) minus five times of contamination (CTN). To compare the Bin Dereplication module with dRep, we also performed dereplication using dRep and assessed the redundancy against standard datasets. Result suggested that dRep can remove part of the redundant bins recovered by co-assembly methods (CA + dRep), while BASALT can efficiently remove all redundant bins (CAB). SA = Single assembly, CA = Co-assembly, CA + dRep = Co-assembly with dereplication using dRep, and CAB = Co-assembly optimized with Bin Dereplication, Refinement and Reassembly modules using BASALT. Datasets processed BASALT only were highlighted with red.

| | | | **Number of non-redundant MAGS in quality score (CPN-5*CTN)** | | | | | | **Number of redundant MAGS in quality score (CPN-5*CTN)** | | | | | | **Redundancy** |
|---|---|---|---|---|---|---|---|---|---|---|---|---|---|---|---|
| | | | ≥90 | ≥80 | ≥70 | ≥60 | ≥50 | Sum | ≥90 | ≥80 | ≥70 | ≥60 | ≥50 | Sum | |
| CAMI-medium | DASTool | SA | 36 | 3 | 5 | 2 | 3 | 49 | 0 | 0 | 0 | 0 | 0 | 0 | 0.0% |
| | | CA | 43 | 5 | 7 | 1 | 2 | 58 | 29 | 9 | 7 | 2 | 2 | 49 | 84.5% |
| | | CA + dRep | 41 | 4 | 5 | 3 | 1 | 54 | 28 | 7 | 2 | 5 | 2 | 45 | 83.3% |
| | VAMB | SA | 23 | 10 | 5 | 3 | 1 | 42 | 0 | 0 | 0 | 0 | 0 | 0 | 0.0% |
| | | CA (Multisplit) | 25 | 13 | 3 | 5 | 3 | 49 | 28 | 3 | 3 | 3 | 1 | 38 | 77.6% |
| | metaWRAP | SA | 36 | 7 | 6 | 5 | 0 | 54 | 0 | 0 | 0 | 0 | 0 | 0 | 0.0% |
| | | CA | 42 | 7 | 5 | 4 | 2 | 60 | 31 | 10 | 5 | 8 | 3 | 57 | 95.0% |
| | | CA + dRep | 40 | 7 | 4 | 4 | 3 | 58 | 24 | 8 | 4 | 7 | 2 | 45 | 77.6% |
| | BASALT | CA | 43 | 8 | 7 | 2 | 1 | 61 | 0 | 0 | 0 | 0 | 0 | 0 | 0.0% |
| CAMI-high | DASTool | SA | 81 | 39 | 18 | 12 | 8 | 158 | 0 | 0 | 0 | 0 | 0 | 0 | 0.0% |
| | | CA | 92 | 42 | 18 | 14 | 12 | 178 | 52 | 36 | 27 | 15 | 17 | 147 | 82.6% |
| | | CA + dRep | 92 | 42 | 17 | 12 | 8 | 171 | 52 | 37 | 26 | 16 | 14 | 145 | 84.8% |
| | | CAB | 104 | 35 | 21 | 10 | 6 | 176 | 0 | 0 | 0 | 0 | 0 | 0 | 0.0% |
| | VAMB | SA | 79 | 37 | 17 | 10 | 12 | 155 | 0 | 0 | 0 | 0 | 0 | 0 | 0.0% |
| | | CA (Multisplit) | 86 | 39 | 20 | 10 | 17 | 172 | 34 | 30 | 16 | 6 | 4 | 90 | 52.3% |
| | | CAB | 85 | 41 | 18 | 12 | 12 | 168 | 0 | 0 | 0 | 0 | 0 | 0 | 0.0% |
| | metaWRAP | SA | 92 | 49 | 41 | 16 | 14 | 212 | 0 | 0 | 0 | 0 | 0 | 0 | 0.0% |
| | | CA | 109 | 52 | 39 | 20 | 18 | 238 | 56 | 60 | 22 | 35 | 19 | 192 | 80.7% |
| | | CA + dRep | 108 | 50 | 40 | 20 | 13 | 231 | 51 | 39 | 23 | 32 | 23 | 168 | 72.7% |
| | | CAB | 112 | 57 | 37 | 13 | 12 | 231 | 0 | 0 | 0 | 0 | 0 | 0 | 0.0% |
| | BASALT | BASALT | 127 | 59 | 37 | 17 | 19 | 259 | 0 | 0 | 0 | 0 | 0 | 0 | 0.0% |

**Table S2.** MAGs used in this study

| MAG ID | BioSample | Taxon CheckM | Reference |
|---|---|---|---|
| bin.T1Sed10.112 | SAMN08575060 | k__Archaea;p__Euryarchaeota | (Vavourakis et al. 2018) |
| bin.CSSed10.265 | SAMN08574522 | k__Archaea;p__Euryarchaeota | |
| bin.B1Sed10.129R1 | SAMN08575184 | k__Archaea;p__Euryarchaeota | |
| bin.T1Sed10.117R1 | SAMN08575063 | k__Archaea;p__Euryarchaeota | |
| bin.T1Sed10.119m | SAMN08575064 | k__Archaea;p__Euryarchaeota | |
| bin.T3Sed10.220R1 | SAMN08574904 | k__Archaea;p__Euryarchaeota | |
| bin.T1Sed10.15 | SAMN08575087 | k__Archaea;p__Euryarchaeota | |
| bin.T1Sed10.130 | SAMN08575071 | k__Archaea (root) | |
| bin.CSSed10.256R1 | SAMN08574517 | k__Archaea (root) | |
| bin.CSSed10.288 | SAMN08574536 | k__Archaea;p__Euryarchaeota | |
| bin.CSSed10.308 | SAMN08574550 | k__Archaea;p__Euryarchaeota | |
| bin.B1Sed10.238R1 | SAMN08575248 | k__Archaea;p__Euryarchaeota | |
| bin.CSSed10.161R1 | SAMN08574455 | k__Archaea;p__Euryarchaeota | |
| bin.B1Sed10.164 | SAMN08575204 | k__Archaea;p__Euryarchaeota | |
| bin.T1Sed10.94 | SAMN08575159 | k__Archaea (root) | |
| bin.T1Sed10.164 | SAMN08575091 | k__Archaea;p__Euryarchaeota | |
| bin.T1Sed10.39R1 | SAMN08575122 | k__Archaea (root) | |
| bin.T1Sed10.101 | SAMN08575053 | k__Archaea;p__Euryarchaeota | |
| bin.CSSed10.238R1 | SAMN08574501 | k__Archaea (root) | |
| bin.B1Sed10.80R1 | SAMN08575294 | k__Archaea;p__Euryarchaeota | |
| bin.CSSed11.322R1 | SAMN08574776 | k__Archaea;p__Euryarchaeota | |
| bin.B1Sed10.198R1 | SAMN08575221 | k__Archaea;p__Euryarchaeota | |
| bin.T3Sed10.350R1 | SAMN08574983 | k__Archaea (root) | |
| bin.B1Sed10.119R1 | SAMN08575176 | k__Archaea;p__Euryarchaeota | |
| bin.CSSed11.243R1 | SAMN08574742 | k__Archaea (root) | |
| bin.T1Sed10.179m | SAMN08575096 | k__Archaea (root) | |
| bin.B1Sed10.49 | SAMN08575271 | k__Bacteria;p__Proteobacteria | |
| bin.T3Sed10.241 | SAMN08574916 | k__Bacteria;p__Bacteroidetes | |
| bin.B1Sed10.40m | SAMN08575263 | k__Bacteria;p__Tenericutes | |
| bin.CSSed10.354m | SAMN08574572 | k__Bacteria (root) | |
| bin.CSSed11.83 | SAMN08574810 | k__Bacteria;p__Thermotogae | |
| bin.T3Sed10.260 | SAMN08574928 | k__Bacteria (root) | |
| bin.T1Sed10.59 | SAMN08575132 | k__Bacteria;p__Proteobacteria | |
| bin.T1Sed10.49 | SAMN08575126 | k__Bacteria;p__Actinobacteria | |
| bin.CSSed10.13m | SAMN08574439 | k__Bacteria;p__Proteobacteria | |
| bin.CSSed10.348R2 | SAMN08574568 | k__Bacteria | |
| bin.CSSed10.131 | SAMN08574441 | k__Bacteria;p__Bacteroidetes | |
| bin.CSSed11.145 | SAMN08574686 | k__Bacteria;p__Planctomycetes | |
| bin.B1Sed10.194 | SAMN08575219 | k__Bacteria;p__Chloroflexi | |
| bin.CSSed11.250 | SAMN08574745 | k__Bacteria;p__Proteobacteria | |
| bin.CSSed10.172R1 | SAMN08574464 | k__Bacteria;p__Bacteroidetes | |

| | | |
|---|---|---|
| bin.T1Sed10.83R1 | SAMN08575149 | k__Bacteria (root) |
| bin.T3Sed10.273 | SAMN08574940 | k__Bacteria;p__Bacteroidetes |
| bin.CSSed11.97R1 | SAMN08574819 | k__Bacteria;p__Planctomycetes |
| bin.CSSed10.289R1 | SAMN08574537 | k__Bacteria;p__Verrucomicrobia |
| bin.B1Sed10.188R1 | SAMN08575216 | k__Bacteria;p__Proteobacteria |
| bin.CSSed11.269R1 | SAMN08574753 | k__Bacteria;p__Chloroflexi |
| bin.CSSed11.114R1 | SAMN08574675 | k__Bacteria (root) |
| bin.T1Sed10.187 | SAMN08575099 | k__Bacteria;p__Bacteroidetes |
| bin.T3Sed10.239 | SAMN08574915 | k__Bacteria;p__Proteobacteria |
| bin.CSSed10.211m | SAMN08574488 | k__Bacteria;p__Tenericutes |
| bin.CSSed10.406 | SAMN08574599 | k__Bacteria;p__Bacteroidetes |
| bin.B1Sed10.85 | SAMN08575298 | k__Bacteria;p__Firmicutes |
| bin.CSSed11.91 | SAMN08574815 | k__Bacteria (root) |
| bin.T3Sed10.96 | SAMN08575031 | k__Bacteria;p__Bacteroidetes |
| bin.T1Sed10.97 | SAMN08575162 | k__Bacteria;p__Planctomycetes |
| bin.T3Sed10.297R1m | SAMN08574949 | k__Bacteria;p__Bacteroidetes |
| bin.T1Sed10.26 | SAMN08575115 | k__Bacteria;p__Firmicutes |
| bin.CSSed10.241R1 | SAMN08574503 | k__Bacteria;p__Firmicutes |
| bin.CSSed10.219R1 | SAMN08574493 | k__Bacteria;p__Verrucomicrobia |
| bin.T3Sed10.182R1 | SAMN08574876 | k__Bacteria;p__Proteobacteria |
| bin.T3Sed10.110R1 | SAMN08574827 | k__Bacteria (root) |
| bin.T3Sed10.12 | SAMN08574833 | k__Bacteria;p__Chloroflexi |
| bin.T1Sed10.85 | SAMN08575151 | k__Bacteria;p__Bacteroidetes |
| bin.T3Sed10.11 | SAMN08574826 | k__Bacteria |
| bin.CSSed11.297R1 | SAMN08574763 | k__Bacteria;p__Proteobacteria |
| bin.T3Sed10.47 | SAMN08575002 | k__Bacteria (root) |
| bin.CSSed11.194 | SAMN08574714 | k__Bacteria;p__Proteobacteria |
| bin.CSSed10.283R1 | SAMN08574533 | k__Bacteria;p__Proteobacteria |
| bin.T3Sed10.170 | SAMN08574867 | k__Bacteria;p__Planctomycetes |
| bin.T3Sed10.218 | SAMN08574902 | k__Bacteria;p__Proteobacteria |
| bin.B1Sed10.114 | SAMN08575174 | k__Bacteria;p__Planctomycetes |
| bin.B1Sed10.223R2 | SAMN08575239 | k__Bacteria (root) |
| bin.CSSed11.41 | SAMN08574783 | k__Bacteria;p__Proteobacteria |
| bin.T3Sed10.264 | SAMN08574931 | k__Bacteria |
| bin.CSSed10.346 | SAMN08574566 | k__Bacteria;p__Spirochaetes |
| bin.CSSed10.9 | SAMN08574660 | k__Bacteria;p__Firmicutes |
| bin.T3Sed10.310R1 | SAMN08574958 | k__Bacteria |
| bin.CSSed11.154 | SAMN08574691 | k__Bacteria;p__Firmicutes |
| bin.B1Sed10.225 | SAMN08575240 | k__Bacteria;p__Tenericutes |
| bin.T1Sed10.69 | SAMN08575138 | k__Bacteria;p__Proteobacteria |
| bin.B1Sed10.220R1 | SAMN08575235 | k__Bacteria;p__Bacteroidetes |
| bin.B1Sed10.223R1 | SAMN08575238 | k__Bacteria (root) |
| bin.B1Sed10.208R1 | SAMN08575227 | k__Bacteria |
| bin.CSSed11.8 | SAMN08574807 | k__Bacteria |

| | | |
|---|---|---|
| bin.T3Sed10.35 | SAMN08574982 | k__Bacteria;p__Proteobacteria |
| bin.CSSed10.305 | SAMN08574549 | k__Bacteria |
| bin.CSSed11.302 | SAMN08574767 | k__Bacteria;p__Proteobacteria |
| bin.T3Sed10.324 | SAMN08574966 | k__Bacteria;p__Proteobacteria |
| bin.B1Sed10.95 | SAMN08575304 | k__Bacteria;p__Proteobacteria |
| bin.CSSed11.246R1m | SAMN08574743 | k__Bacteria;p__Proteobacteria |
| bin.CSSed10.82m | SAMN08574653 | k__Bacteria;p__Cyanobacteria |
| bin.T3Sed10.329R1 | SAMN08574969 | k__Bacteria |
| bin.B1Sed10.159 | SAMN08575200 | k__Bacteria;p__Firmicutes |
| bin.CSSed10.295 | SAMN08574541 | k__Bacteria;p__Verrucomicrobia |
| bin.CSSed11.197 | SAMN08574715 | k__Bacteria;p__Chloroflexi |
| bin.B1Sed10.66 | SAMN08575284 | k__Bacteria;p__Proteobacteria |
| bin.CSSed11.191 | SAMN08574713 | k__Bacteria;p__Proteobacteria |
| bin.T3Sed10.257 | SAMN08574925 | k__Bacteria;p__Spirochaetes |
| bin.CSSed11.109 | SAMN08574673 | k__Bacteria;p__Firmicutes |
| bin.CSSed10.184 | SAMN08574474 | k__Bacteria;p__Verrucomicrobia |
| bin.T1Sed10.77m | SAMN08575145 | k__Bacteria;p__Firmicutes |
| bin.CSSed11.242R1 | SAMN08574741 | k__Bacteria;p__Proteobacteria |
| bin.CSSed11.54R1 | SAMN08574792 | k__Bacteria;p__Bacteroidetes |
| bin.CSSed10.104 | SAMN08574421 | k__Bacteria (root) |
| bin.CSSed10.216R1 | SAMN08574491 | k__Bacteria |
| bin.T3Sed10.347R1 | SAMN08574979 | k__Bacteria |
| bin.CSSed10.109m | SAMN08574424 | k__Bacteria;p__Tenericutes |
| bin.B1Sed10.179 | SAMN08575211 | k__Bacteria;p__Proteobacteria |
| bin.CSSed10.181R1 | SAMN08574472 | k__Bacteria;p__Spirochaetes |
| bin.T3Sed10.262 | SAMN08574930 | k__Bacteria;p__Bacteroidetes |
| bin.CSSed11.258 | SAMN08574748 | k__Bacteria;p__Planctomycetes |
| bin.T1Sed10.160 | SAMN08575090 | k__Bacteria;p__Firmicutes |
| bin.T3Sed10.326m | SAMN08574967 | k__Bacteria (root) |
| bin.CSSed11.31 | SAMN08574770 | k__Bacteria;p__Bacteroidetes |
| bin.CSSed10.472R1 | SAMN08574626 | k__Bacteria (root) |
| bin.CSSed10.375 | SAMN08574578 | k__Bacteria;p__Chloroflexi |
| bin.CSSed10.237R1 | SAMN08574499 | k__Bacteria;p__Firmicutes |
| bin.T3Sed10.22m | SAMN08574903 | k__Bacteria;p__Deinococcus-Thermus |
| bin.T1Sed10.3 | SAMN08575118 | k__Bacteria;p__Proteobacteria |
| bin.CSSed10.361 | SAMN08574575 | k__Bacteria;p__Proteobacteria |
| bin.T1Sed10.134 | SAMN08575075 | k__Bacteria (root) |
| bin.CSSed11.96 | SAMN08574818 | k__Bacteria |
| bin.CSSed10.133 | SAMN08574442 | k__Bacteria;p__Firmicutes |
| bin.CSSed10.186 | SAMN08574475 | k__Bacteria (root) |
| bin.T3Sed10.202R1 | SAMN08574890 | k__Bacteria (root) |
| bin.CSSed10.434R1 | SAMN08574612 | k__Bacteria;p__Firmicutes |
| bin.CSSed11.315 | SAMN08574772 | k__Bacteria (root) |

| | | |
|---|---|---|
| bin.CSSed10.474R1 | SAMN08574627 | k__Bacteria;p__Proteobacteria |
| bin.CSSed11.173R1 | SAMN08574702 | k__Bacteria (root) |
| bin.T3Sed10.304 | SAMN08574956 | k__Bacteria;p__Cyanobacteria |
| bin.CSSed11.288R1 | SAMN08574759 | k__Bacteria;p__Actinobacteria |
| bin.CSSed10.297R1 | SAMN08574542 | k__Bacteria (root) |
| bin.T1Sed10.198m | SAMN08575103 | k__Bacteria (root) |
| bin.CSSed11.312R1 | SAMN08574771 | k__Bacteria;p__Planctomycetes |
| bin.CSSed10.89 | SAMN08574659 | k__Bacteria;p__Proteobacteria |
| bin.CSSed10.77 | SAMN08574651 | k__Bacteria;p__Bacteroidetes |
| bin.CSSed10.448R1 | SAMN08574619 | k__Bacteria;p__Proteobacteria |
| bin.CSSed10.205R1 | SAMN08574485 | k__Bacteria (root) |
| bin.T3Sed10.256R1 | SAMN08574923 | k__Bacteria;p__Verrucomicrobia |
| bin.B1Sed10.231 | SAMN08575245 | k__Bacteria;p__Planctomycetes |
| bin.T3Sed10.271R1 | SAMN08574939 | k__Bacteria;p__Proteobacteria |
| bin.CSSed11.263 | SAMN08574751 | k__Bacteria (root) |
| bin.CSSed11.174 | SAMN08574703 | k__Bacteria |
| bin.CSSed11.304 | SAMN08574769 | k__Bacteria;p__Proteobacteria |
| bin.CSSed11.155 | SAMN08574692 | k__Bacteria;p__Proteobacteria |
| bin.B1Sed10.227R1 | SAMN08575242 | k__Bacteria;p__Firmicutes |
| bin.CSSed10.402R1 | SAMN08574597 | k__Bacteria (root) |
| bin.T3Sed10.236R1 | SAMN08574914 | k__Bacteria |
| bin.CSSed11.170R1 | SAMN08574698 | k__Bacteria;p__Chloroflexi |
| bin.CSSed10.440 | SAMN08574616 | k__Bacteria (root) |
| bin.T3Sed10.354 | SAMN08574984 | k__Bacteria (root) |
| bin.T3Sed10.102R1 | SAMN08574823 | k__Bacteria (root) |
| bin.CSSed10.113R1 | SAMN08574428 | k__Bacteria (root) |
| bin.CSSed10.128 | SAMN08574438 | k__Bacteria;p__Proteobacteria |
| bin.CSSed10.310 | SAMN08574552 | k__Bacteria (root) |
| bin.CSSed10.457 | SAMN08574621 | k__Bacteria (root) |
| bin.B1Sed10.118 | SAMN08575175 | k__Bacteria;p__Planctomycetes |
| bin.T3Sed10.319 | SAMN08574963 | k__Bacteria;p__Verrucomicrobia |
| bin.B1Sed10.209R1 | SAMN08575229 | k__Bacteria;p__Planctomycetes |
| bin.T3Sed10.223R1 | SAMN08574905 | k__Bacteria;p__Proteobacteria |
| bin.CSSed11.318R1 | SAMN08574773 | k__Bacteria;p__Chloroflexi |
| bin.CSSed10.381R1 | SAMN08574584 | k__Bacteria (root) |
| bin.CSSed10.419R1 | SAMN08574608 | k__Bacteria;p__Planctomycetes |
| bin.CSSed10.249R1 | SAMN08574510 | k__Bacteria;p__Proteobacteria |
| bin.CSSed11.257m | SAMN08574747 | k__Bacteria (root) |
| bin.B1Sed10.221R1 | SAMN08575236 | k__Bacteria;p__Planctomycetes |
| bin.CSSed10.423 | SAMN08574609 | k__Bacteria |
| bin.T3Sed10.344R1 | SAMN08574978 | k__Bacteria;p__Cyanobacteria |
| bin.CSSed11.89 | SAMN08574813 | k__Bacteria |
| bin.T3Sed10.224 | SAMN08574906 | k__Bacteria;p__Verrucomicrobia |
| bin.CSSed10.250 | SAMN08574511 | k__Bacteria (root) |

| | | | |
|---|---|---|---|
| bin.CSSed10.428R1 | SAMN08574610 | k__Bacteria;p__Firmicutes | |
| bin.T3Sed10.376R1m | SAMN08574996 | k__Bacteria;p__Cyanobacteria | |
| bin.CSSed10.463m | SAMN08574622 | k__Bacteria;p__Cyanobacteria | |
| bin.CSSed10.43 | SAMN08574611 | k__Bacteria;p__Firmicutes | |
| bin.CSSed10.466R1 | SAMN08574623 | k__Bacteria (root) | |
| TACK_SB0675_bin_21 | SAMN12598637 | k__Archaea;p__Crenarchaeota | (Engelberts et al. 2020) |
| TACK SB0662_bin_33 | SAMN12598268 | k__Archaea;p__Crenarchaeota | |
| LC_3 | SAMN04924817 | k__Archaea;p__Heimdallarchaeota | (Zaremba-Niedzwiedzka et al. 2017) |
| AB_125 | SAMN04924818 | k__Archaea;p__Heimdallarchaeota | |
| LC_2 | SAMN04924815 | k__Archaea;p__Heimdallarchaeota | |
| LCB_4 | SAMN04924820 | k__Archaea;p__Odinarchaeota | |
| SZ_4_bin3.344 | SAMN10218971 | k__Archaea;p__Thorarchaeota | (Cai et al. 2020) |
| SZ_4_bin10.384 | SAMN10219203 | k__Archaea;p__Helarchaeota | |
| SZ_4_bin8.338 | SAMN10218974 | k__Archaea;p__Lokiarchaeota | |
| DZG_bin1.240 | SAMN10218969 | k__Archaea;p__Lokiarchaeota | |
| MP8T_1 | SAMN06640692 | k__Archaea;p__Thorarchaeota | (Zhou et al. 2019) |
| MP11T_1 | SAMN06640694 | k__Archaea;p__Thorarchaeota | |
| Hel_GB_B | SAMN09406172 | k__Archaea;p__Helarchaeota | (Seitz et al. 2019) |
| Hel_GB_A | SAMN09406154 | k__Archaea;p__Helarchaeota | |
| Helarchaeota_1 | SAMN14414633 | k__Archaea;p__Helarchaeota | |
| Helarchaeota_2 | SAMN14414634 | k__Archaea;p__Helarchaeota | |
| B53_G9 | SAMN09215251 | k__Archaea;p__Lokiarchaeota | |
| AS27yjCOA_147 | SAMN13894882 | k__Archaea;p__Lokiarchaeota | (Campanaro et al. 2020) |
| CR_4 | SAMN04958229 | k__Archaea;p__Lokiarchaeota | (Kauffman et al. 2018) |
| MK-D1 | SAMN12405820 | k__Archaea;p__Lokiarchaeota | (Imachi et al. 2020) |
| Bin_342 | SAMN13151233 | k__Archaea;p__Lokiarchaeota | (Wong et al. 2020) |
| Bin_186 | SAMN13151198 | k__Archaea;p__Helarchaeota | |
| Bin_229 | SAMN13151180 | k__Archaea;p__Lokiarchaeota | |
| Bin_204 | SAMN13151185 | k__Archaea;p__Lokiarchaeota | |
| RBin_035 | SAMN13151185 | k__Archaea;p__Lokiarchaeota | |
| RBin_125 | SAMN13151222 | k__Archaea;p__Lokiarchaeota | |
| Bin_485 | SAMN13151252 | k__Archaea;p__Lokiarchaeota | |
| AMARA_1S | SAMN10768446 | k__Archaea;p__Lokiarchaeota | (Bulzu et al. 2019) |
| AMARA_7 | SAMN10768449 | k__Archaea;p__Lokiarchaeota | |
| TEKIR_21 | SAMN10768457 | k__Archaea;p__Lokiarchaeota | |
| AMARA_5S | SAMN10768448 | k__Archaea;p__Lokiarchaeota | |
| Loki_b32 | SAMN07236659 | k__Archaea;p__Lokiarchaeota | PRJNA383916 |
| Loki_b31 | SAMN07236658 | k__Archaea;p__Lokiarchaeota | |
| Hel_238_bin13 | SAMN14451652 | k__Archaea;p__Lokiarchaeota | (Yin et al. 2020) |
| Hel_238_bin90 | SAMN14451653 | k__Archaea;p__Lokiarchaeota | |
| Hel_238_bin105 | SAMN14451654 | k__Archaea;p__Lokiarchaeota | |
